# Flexynesis: A deep learning framework for bulk multi-omics data integration for precision oncology and beyond

**DOI:** 10.1101/2024.07.16.603606

**Authors:** Bora Uyar, Taras Savchyn, Ricardo Wurmus, Ahmet Sarigun, Mohammed Maqsood Shaik, Vedran Franke, Altuna Akalin

## Abstract

Accurate decision making in precision oncology depends on integration of multimodal molecular information, such as the genetic data, gene expression, protein abundance, and epigenetic measurements. Deep learning methods facilitate integration of heterogeneous datasets. However, almost all published deep learning-based bulk multi-omics integration methods have constrained usability. They suffer from lack of transparency, modularity, deployability, and are applicable exclusively to narrow tasks. To address these limitations, we introduce Flexynesis, a versatile tool designed with usability, and adaptability in mind. Flexynesis streamlines data processing, enforces structured data splitting, and ensures rigorous model evaluation. It offers unsupervised feature selection, different omics layer fusion options, and hyperparameter tuning. Users can choose from distinct architectures – fully connected networks, variational autoencoders, multi-triplet networks, graph neural networks, and cross-modality encoding networks. Each model is complemented with a straightforward input interface and standardized training, evaluation, and feature importance quantification methods, enabling easy incorporation into data integration pipelines. For improved user experience, Flexynesis supports features such as on-the-fly task determination and compatibility with regression, classification, and survival modeling. It accommodates multi-task prediction of a mixture of numerical/categorical outcome variables with a tolerance for missing labels. We also developed an extensive benchmarking pipeline, showcasing the tool’s capability across diverse real-life datasets. This toolset should make deep-learning based bulk multi-omics data integration in the context of clinical/pre-clinical data analysis and marker discovery more accessible to a wider audience with or without experience in deep-learning development. Flexynesis is available at https://github.com/BIMSBbioinfo/flexynesis and can be installed from https://pypi.org/project/flexynesis/.

**Graphical Abstract:**
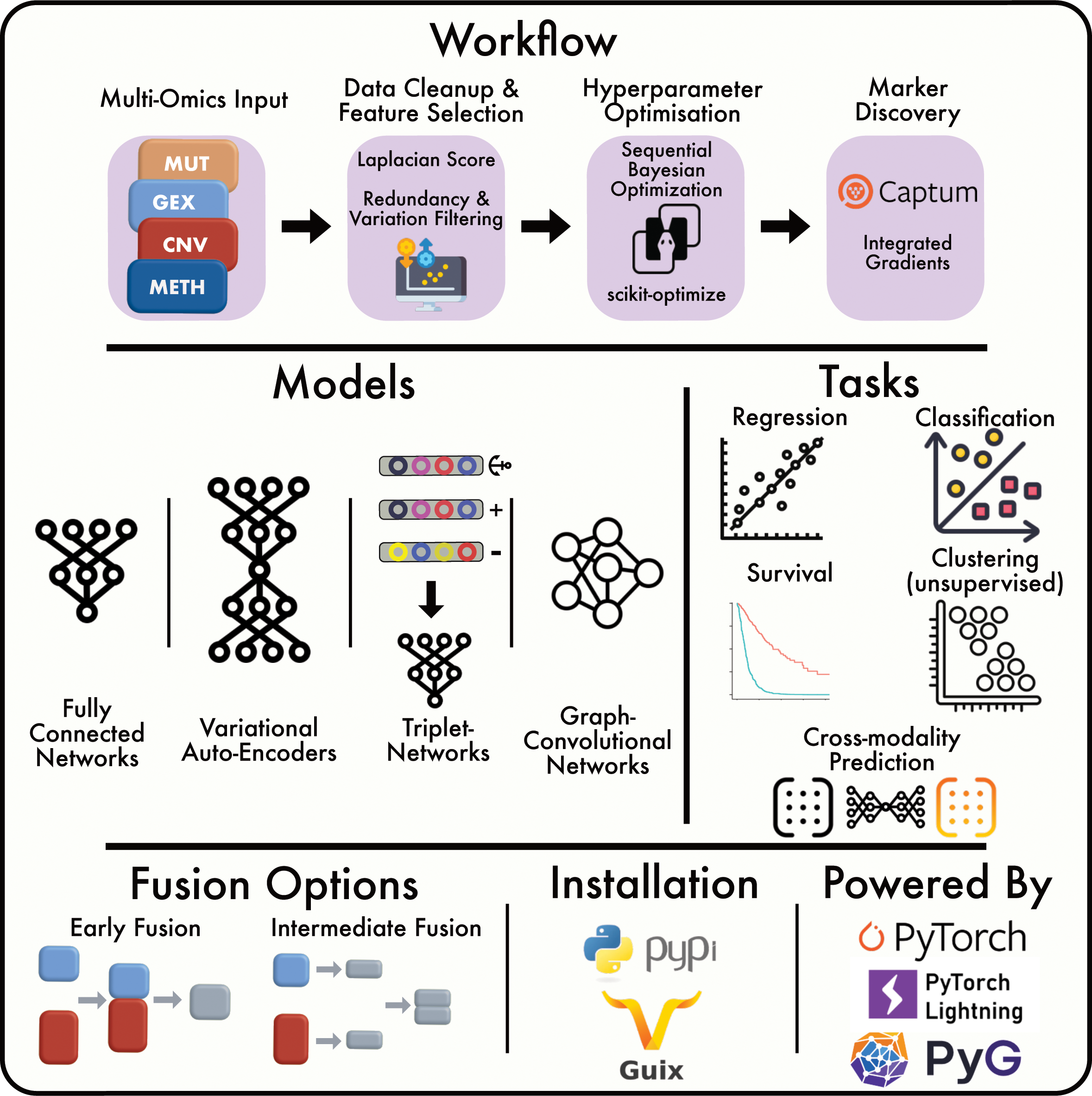
Summary of the Flexynesis data integration and analysis workflow.

## Introduction

Cancer is a complex disease primarily resulting from genomic aberrations. The disease is marked by abnormal cell growth, invasive proliferation, and tissue malfunction, impacting twenty million individuals and causing ten million yearly deaths worldwide [1]. To bypass protective mechanisms, cancer cells must acquire several key characteristics, such as resistance to cell death, immune evasion, tissue invasion, growth suppressor evasion, and sustained proliferative signaling [2]. Unlike rare genetic disorders, caused by few genetic variations, complex diseases, like cancer, require a comprehensive understanding of interactions between various cellular regulatory layers. This entails data integration from various omics layers, such as the transcriptome, epigenome, proteome, genome, metabolome, and microbiome [3]. In clinical settings, genome-informed diagnostics to identify disease-causing variants are already in use [4]. However, capturing the complexity of most cancers requires more than a panel of genomic markers. Multi-omics profiling is a vital step toward understanding not only cancer but other complex diseases like cardiovascular and neurological disorders [5–7]. Proof-of-concept studies have shown the benefits of multi-omics patient profiling for health monitoring, treatment decisions, and knowledge discovery [8]. Recent longitudinal clinical studies in cancer are evaluating the effects of multi-omics-informed clinical decisions compared to standard of care [9]. Addressing this need for multi-omic profiling to improve the understanding of complex diseases, major international initiatives have developed multi-omic databases such as The Cancer Genome Atlas (TCGA), the Cancer Cell Line Encyclopedia (CCLE) [10] to enhance molecular profiling of tumors and disease models.

While cell regulation at the molecular level is highly interconnected, redundant, and has non-linear relationships between components, the information about these intricate relationships is usually isolated in different molecular data modalities. Each molecular profile is measured one assay at a time (as in assays developed for profiling the transcriptome, the genome, the methylome etc), however, all the different layers of molecular information are in actuality in a cross-talk with one another. Therefore, it is important to capture the non-linear relationships, and impacts of disruptions of the different components of the cellular machinery by combining the disparate data modalities into a more meaningful synthesis. However, the high dimensionality of molecular assays and heterogeneity of the studied diseases create computational challenges.

The challenges of multi-omics data integration prompted development of various machine learning algorithms, including deep learning approaches [11,12]. Available benchmarking studies that compared different deep-learning-based methods for multi-omics integration for classification and regression tasks [13,14] have shown that none of the methods clearly outperformed others in all the tasks at hand. This necessitates a flexible and reproducible approach that provides adaptable architectures for solving each computational task.

Before setting out to develop yet another deep learning-based multi-omics integration method, despite the availability of the myriad of published studies [11], we collated a survey of available bulk multi-omics data integration methods to see which tools can be easily adapted for our own translational research projects (Supplementary Table 1). Such projects usually include heterogeneous cohorts of cancer patients and pre-clinical disease models with multi-omics profiles. A primary issue we observed with existing methods is their limited reusability or adaptability to different datasets and contexts. The majority of published approaches do not provide accompanying code, severely limiting their accessibility and applicability. Even when the code is available, it often exists as an unpackaged collection of scripts or notebooks. Such a disorganized format makes these methods difficult, if not impossible, to install, reuse, and incorporate into existing bioinformatics pipelines. Out of the 80 studies collated, 29 studies provide no codebase. 45 studies provide collections of scripts/notebooks, with the goal of reproducing the findings in the published study rather than serving as a generic tool for multi-omics integration. While these methods (Supplementary Table 1) are valuable contributions to the scientific community, they still require extensive customization to make them usable for different datasets and tasks.

Besides lacking readily available code, published methods suffer from one or more of the criteria that are crucial for ensuring the reliability and reproducibility of machine learning applications. Standard operating procedures such as training/validation/test splits, hyperparameter optimization, feature selection, and marker discovery are frequently overlooked or manually defined, without any accompanying documentation again underscoring the arduous amount of work needed to adapt these approaches for custom problems.

Another limitation of current deep learning methods is their narrow task specificity. Many tools are designed exclusively for specific applications, such as regression, survival modeling, or classification. Comprehensive multi-omics data analysis frequently requires a mixture of such tasks, however, the specialization of already existing tools restricts their applicability.

While deep learning methods are sometimes considered as superior, classical machine learning algorithms frequently outperform them [15–17]. This performance differential is not immediately apparent, and often not tested with the currently existing tools, requiring users to undertake extensive benchmarking to uncover the most effective solution to their specific problem.

Addressing these challenges, we introduce Flexynesis, a novel deep learning framework designed to overcome the above-mentioned limitations. We demonstrate the versatility of Flexynesis through various use cases, including drug response prediction, cancer subtype modeling, survival analysis, and biomarker discovery. We demonstrate how to handle multiple tasks simultaneously, supporting a combination of regression, classification, and survival tasks. We show use-cases where the flexibility of neural networks can be utilized in different prediction tasks by building models of both unsupervised and supervised tasks, with one or more supervision heads, and symmetric (auto-encoders) and asymmetric (cross-modality) encoders.

To further enhance its utility, we provide an accessory pipeline and a collection of datasets for benchmarking different flavors of Flexynesis. This benchmarking includes a comparison to classical machine learning methods (Random Forests, Support Vector Machines, and Random Survival Forests).

In summary, the landscape of published deep learning methods for bulk multi-omics data integration is fraught with challenges that hinder their effective reuse and integration into broader bioinformatics workflows. This manuscript addresses these challenges and introduces Flexynesis, a comprehensive solution designed to enhance the utility and applicability of deep learning in multi-omics data analysis.

## Results

We designed Flexynesis for automated construction of predictive models of one or more outcome variables. For each outcome variable, a supervisor multi-layer-perceptron (MLP) is attached onto the encoder networks (a selection of fully connected or graph-convolutional encoders) to perform the modeling task. Clinically relevant machine learning tasks such as drug response prediction (regression), disease subtype prediction (classification), and survival modeling (right-censored regression) tasks are all possible as individual variables or as a mixture of variables, such that each outcome variable has an impact on the low-dimensional sample embeddings (latent variables) derived from the encoding networks.

### Single-task modeling : Predicting only one outcome variable

In Figure 1, we demonstrate the different kinds of modeling tasks that are possible with Flexynesis using a single outcome variable (single MLP) as regression (Figure 1A), classification (Figure 1B), and survival models (Figure 1C). For the regression task, we trained Flexynesis on multi-omics (gene expression and copy-number-variation) data from cell lines from the CCLE database [10] to predict the cell line sensitivity levels to the drugs Lapatinib, a tyrosine kinase inhibitor, and Selumetinib, a MEK inhibitor. We evaluated the performance of the trained model on the cell lines from the GDSC2 database [18] which were also treated with the same drugs, where we observed a high correlation between the known drug response values and the predicted response values for both drugs (Figure 1A).

**Figure 1:**
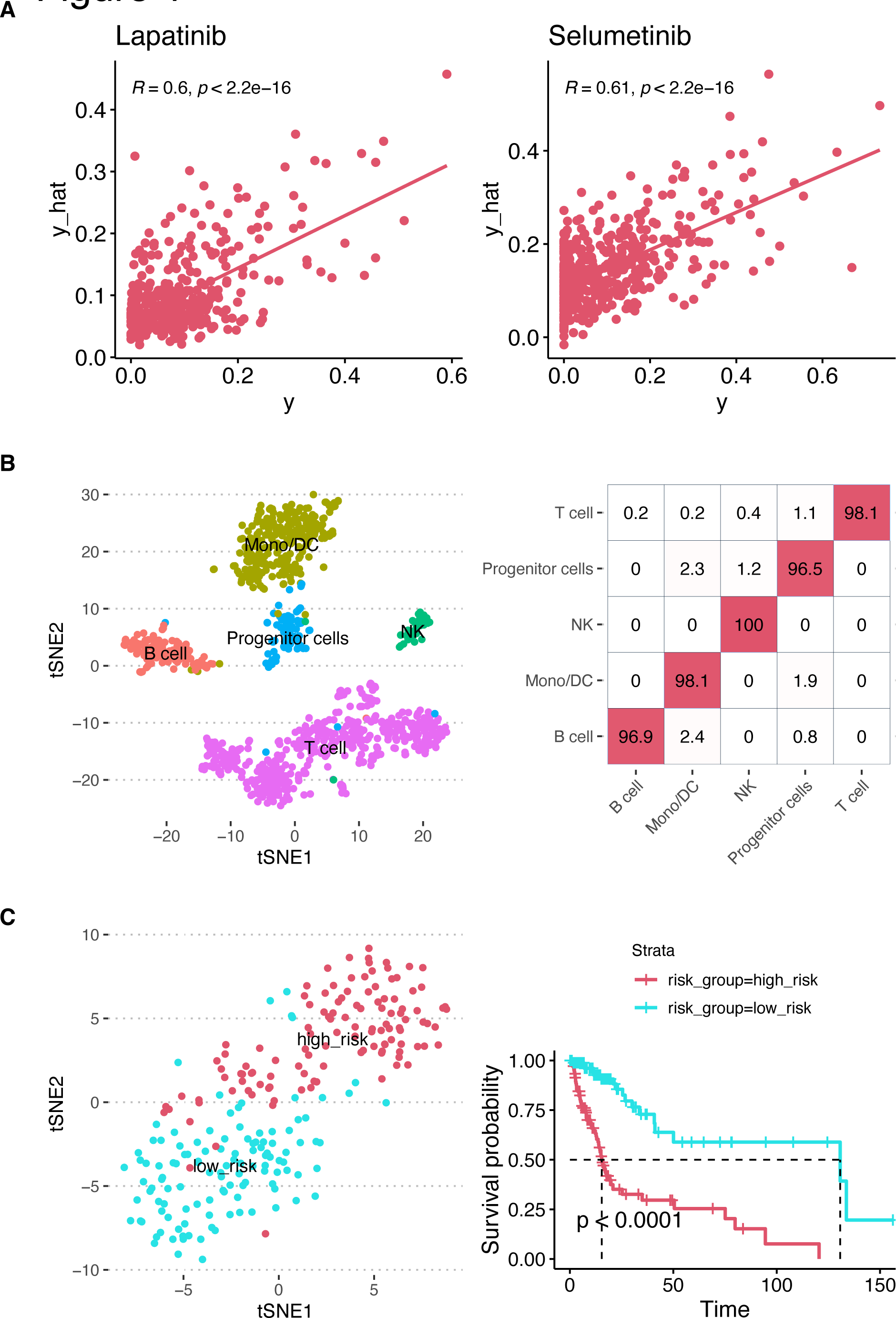
Flexynesis supports single-task modeling for regression (panel A), classification (panel B), and survival (panel C). For all three tasks, both a fully-connected-network and a supervised variational auto-encoder was trained and best-performing model’s results were presented. **A)** performance evaluation of Flexynesis on drug response prediction of a model trained on 1051 cell lines from CCLE (using RNA and CNV profiles) and evaluated on 1075 cell lines from GDSC2 for the drugs Lapatinib and Selumetinib. The x-axis depicts observed drug response values (AAC-recomputed as in Pharmacogx package [55]) and the y-axis depicts the predicted drug response values for the test samples. **B)** evaluation of Flexynesis on a cell type classification problem, where the model is trained on 5000 bone marrow cells with CITE-seq data (using RNA and ADT profiles) and evaluated on 5000 cells from the same dataset. The t-SNE plot represents the sample embeddings obtained from the model encoder and the heatmap represents the confusion matrix for the known and predicted cell type labels for the test samples. **C)** evaluation of Flexynesis on a survival modeling task on a merged cohort of LGG and GBM patient samples (using mutations and copy-number-alteration profiles). The model is trained on 557 samples and evaluated on 239 test samples. The tSNE plot depicts the sample embeddings obtained from the model encoder for the test samples colored by the predicted Cox proportional hazard risk scores stratified into “high-risk” and “low-risk” based on the median risk score. The Kaplan-Meier-Plot represents the survival stratification of the test samples based on this risk stratification (p-value < 0.0001, log-rank test).

For the single-variable classification task, we chose a single-cell multi-omics dataset (CITE-Seq) of PBMCs [19], where we built a cell type label prediction model on 5000 cells using gene expression and antibody-derived tags (ADT) as the input data modalities and evaluated the model performance on the hold-out dataset of 5000 cells, which resulted in a high accuracy cell type label classifier as reflected by the sample embeddings of the cells from the holdout dataset and the confusion-matrix of known and predicted cell type labels (Figure 1B).

As the third type of modeling task, we demonstrate survival modeling using Flexynesis on a combined cohort of lower grade glioma (LGG) and glioblastoma multiforme (GBM) patient samples [20]. For survival modeling, a supervisor MLP with Cox Proportional Hazards loss function is used to guide the network to learn patient-specific risk scores based on the input overall survival endpoints as has been demonstrated previously [21]. After training the model on 70% of the samples, we predicted the risk scores of the remaining test samples (30%) and split the risk scores by the median risk value in the cohort. The embeddings visualized based on the median risk score stratification shows that the test samples are clearly separable in the sample embedding space, which is also confirmed by the Kaplan-Meier survival plot, which shows a significant separation of patients in terms of predicted risk scores (Figure 1C).

### Multi-task Modeling: Joint prediction of multiple outcome variables

While being able to build deep learning models with any of the regression/classification/survival tasks individually offers an improved user experience, this is also usually possible with classical machine learning methods. The actual flexibility of deep learning is more evident in a multi-task setting where more than one MLPs are attached on top of the sample encoding networks, thus the embedding space can be shaped by multiple clinically relevant variables. This flexibility is even more pronounced in the presence of missing labels for one or more of the variables, which is tolerated by Flexynesis.

To demonstrate the use of multi-task modeling, we trained models on 70% of the METABRIC dataset (a metastatic breast cancer cohort with multi-omics profiles of 2509 patients) [22] and obtained the embeddings for the 30% of the samples. In order to compare and contrast the effect of multi-task modeling with single-task modeling, we chose two clinically relevant variables for this cohort: subtype labels (CLAUDIN_SUBTYPE) and chemotherapy treatment status (CHEMOTHERAPY). We built three different models: a single-task model using only the subtype labels (Figure 2A), a single-task model using only the chemotherapy status of the patients (Figure 2B), and finally a multi-task model using both subtype labels and chemotherapy status as outcome variables (Figure 2C). Coloring the test samples by the subtype labels and chemotherapy status, we can observe that the sample embeddings obtained exclusively for the subtype modeling reflect a clear clustering of samples by subtype, but not by the chemotherapy status (Figure 2A). Similarly, the sample embeddings obtained from the model trained exclusively with the chemotherapy status as outcome variable shows a clear separation of samples by treatment status, however the separation by subtypes is not as evident anymore (Figure 2B). In the multi-task setting where the model had two MLPs (one for subtype labels and one for chemotherapy status), the sample embeddings show a clear separation of both by the subtype labels and also the chemotherapy status (Figure 2C).

**Figure 2:**
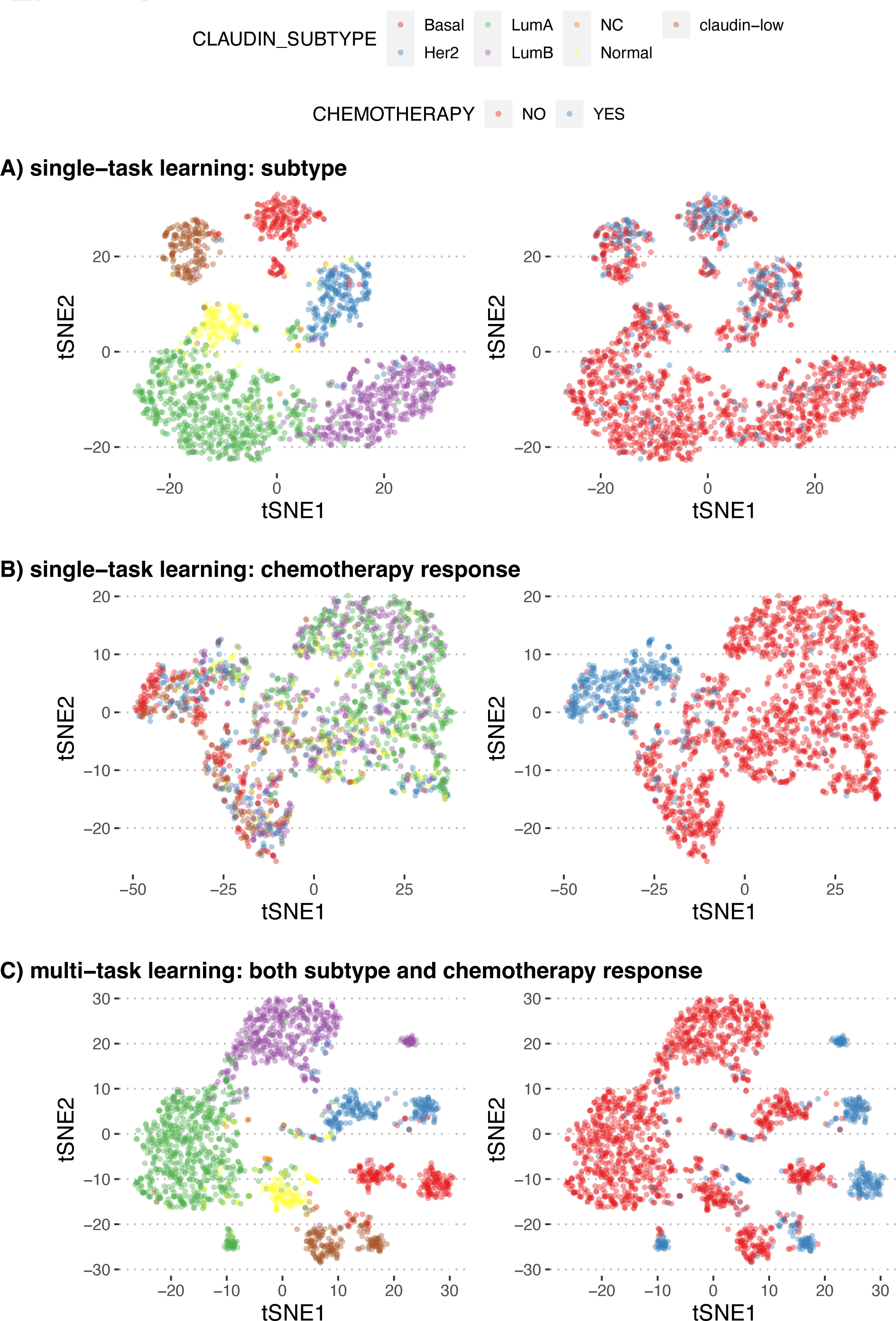
t-SNE plots representing sample embeddings from 561 test samples of metastatic breast cancer from the METABRIC study. These plots compare the impact of single-task and multi-task modeling on the clustering of samples by clinical variables. Plots on the left are colored by the breast cancer subtype and the plots on the right are colored by the treatment status. **A)** Single-Task Model – Breast Cancer Subtypes: t-SNE visualization of test sample embeddings obtained from a single-task model trained exclusively to predict breast cancer subtypes. **B)** Single-Task Model – Chemotherapy Status: t-SNE plot visualization of test sample embeddings from a model trained only to predict the chemotherapy status of patients, showing the segregation capability of the single-task model with respect to treatment status. **C)** Multi-Task Model – Subtypes and Chemotherapy Status: t-SNE plot of test sample embeddings from a multi-task model trained with dual supervisor heads: one for breast cancer subtypes and another for chemotherapy status. The plot shows how multi-task learning influences the embedding space, enhancing the separation of samples based on both clinical variables simultaneously.

We also analyzed the LGG and GBM cohort (from Figure 1C) in a multi-task setting where we attached three separate MLPs on the encoder layers: a regressor to predict the patient’s age (AGE), a classifier to predict the histological subtype (HISTOLOGICAL DIAGNOSIS), and another survival head to model the survival outcomes of the patients (OS_STATUS). Concurrently training the model with three different tasks at the same time, we inspected the sample embeddings and observed that older patients with high risk scores have the glioblastoma subtype, while younger patients with lower risk scores have the other subtypes, where low risk young patients can still be distinguished mainly by histological subtype (Figure 3A). Thus, training the model on three clinically relevant variables helps us obtain sample embeddings that reflect all three variables in a hierarchical manner. Inspecting the top markers for each of these variables, we observe common genes for all three variables such as IDH1, IDH2, ATRX, PIK3CA, and EGFR (Figure 3B), which could be explained by the fact that the clinical variables such as age and histological subtype are correlated with the survival outcomes of the patients, underpinning the importance of these genes in the etiology of the gliomas, which have been extensively studied and reported before [23].

**Figure 3:**
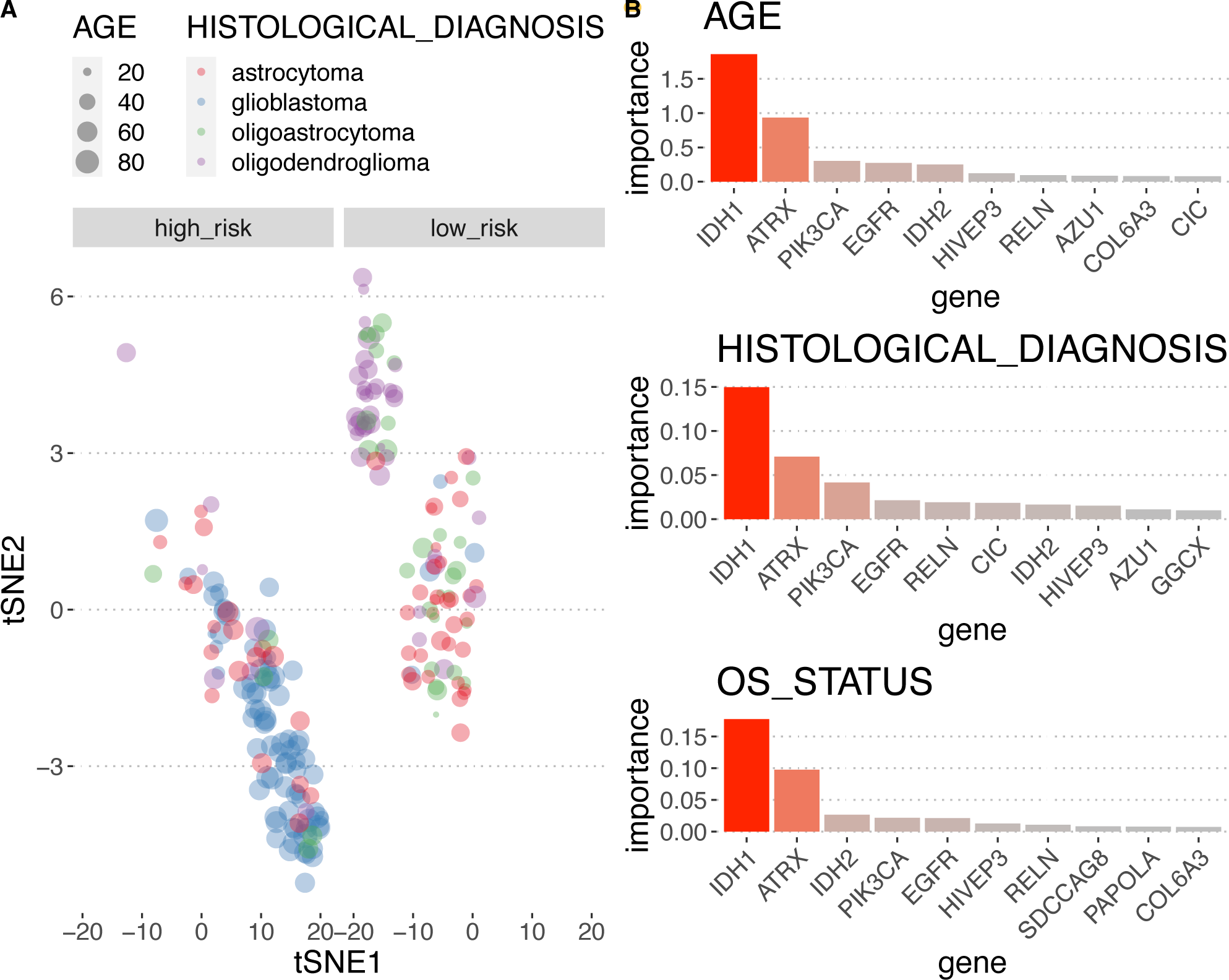
Flexynesis can be trained concurrently for all three types of tasks: regression, classification, and survival at a single run. The model was trained on 557 training samples from the merged cohort of the LGG and GBM patient samples with three supervisor heads: a regressor for the patient age (AGE), a classifier for the histological diagnosis, and a survival head for the overall survival status of the patient (OS_STATUS). Panel **A)** displays the tSNE visualization of the sample embeddings for 239 test samples, where the size of the points reflect the age of the patient, the colors represent the histological diagnosis, and the samples were stratified into high-risk and low-risk groups based on the predicted risk scores for each patient. The sample embeddings reflect the impact of all three clinical variables concurrently. Panel **B)** displays the top 10 most important features discovered for each supervisor head for the patient’s age, histological subtype, and survival status.

### Unsupervised learning: finding groups and general patterns

One of the main architectures provided in Flexynesis is the variational auto-encoders (VAE) with maximum mean discrepancy (MMD) loss [24]. While VAEs are usually employed in unsupervised training tasks, in Flexynesis they can be used for both supervised and unsupervised tasks. In the absence of any target outcome variables (in other words, without any additional MLP modules attached on top of the encoders), the network behaves as a VAE-MMD where the sole goal is to reconstruct the input data matrices, while generating embeddings that follow a Gaussian distribution due to the MMD loss.

As a proof of principle experiment, we trained a VAE-MMD model without any attached supervisor MLPs, to test the unsupervised dimension reduction capabilities on 21 cancer types from the TCGA resource using gene expression and methylation as input modalities. Applying k-means clustering (k from 18 to 24), we obtained a clustering of the samples based on the trained sample embeddings. The tSNE representation of the resulting sample embeddings shows a clear separation of unsupervised clusters (Figure 4A) and the known sample labels (Figure 4B) with a good correspondence between unsupervised clusters and known sample labels (adjusted mutual information: 0.78) (Figure 4C).

**Figure 4:**
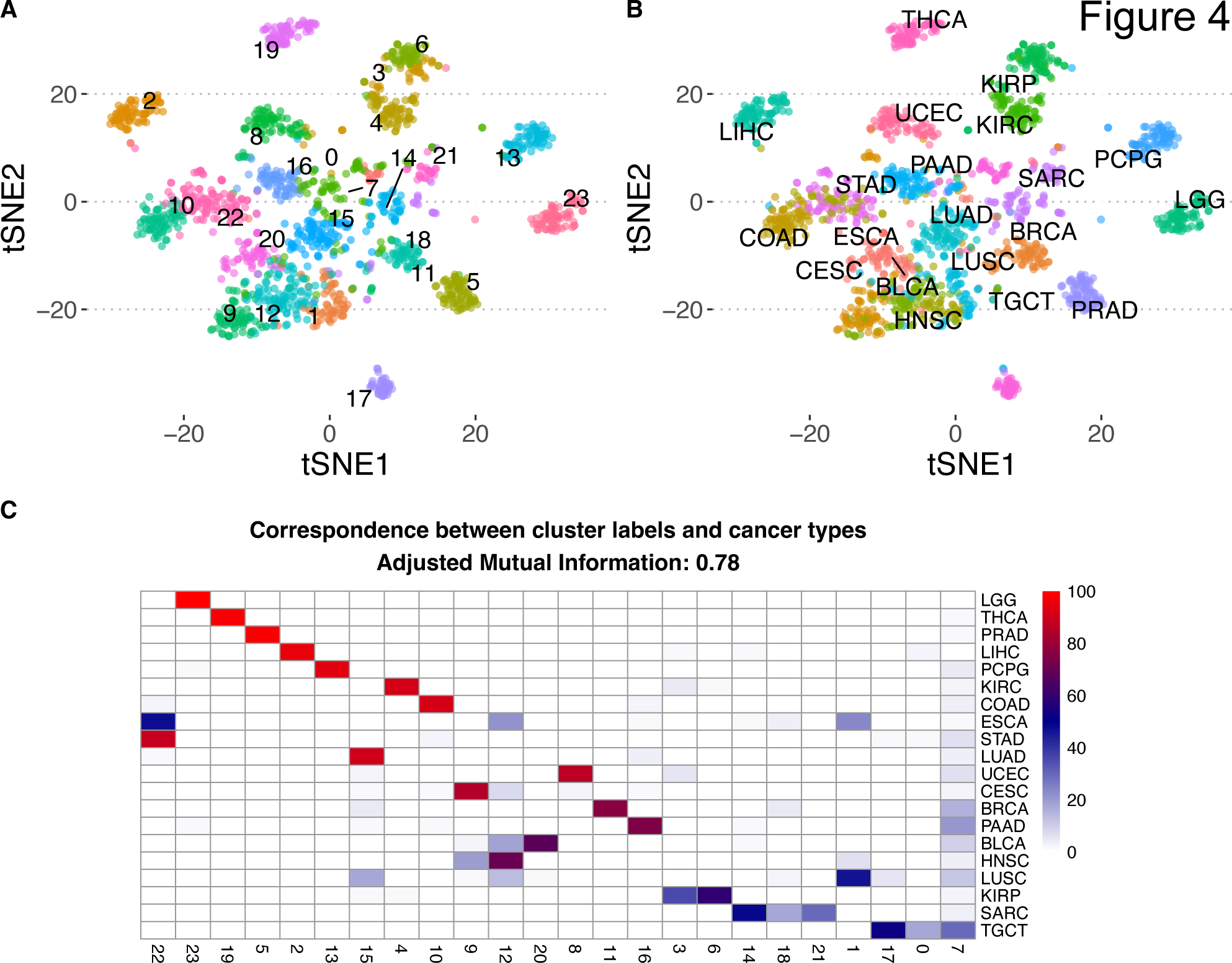
Flexynesis can be used for unsupervised training and clustering. The figure displays the unsupervised analysis of 21 cancer types from the TCGA study for 1600 samples (80 samples per cancer type were randomly selected). Panel **A)** displays the tSNE plot of the training sample embeddings colored by the best performing clustering scheme using the k-means algorithm for values of 18<=k<=24, where best clustering was selected by the best silhouette score (k = 24). Panel **B)** the same tSNE plot as in A) but colored by the known cancer type labels. Panel **C)** The river plot displays the concordance between the cluster labels from panel A) and known cancer type labels from panel B), where the adjusted mutual information score is 0.78.

### Cross-modality learning: Transferring knowledge between different omic data types

While variational autoencoders are designed to reconstruct the initial input data, this can be formulated in a different fashion such that the goal of the reconstruction is a set of matrices different from the inputs. Thus, it is possible to build models where the input data modalities differ from the output data modalities. For instance, a gene expression data matrix could be used to reconstruct a mutation data matrix, thus learning how to translate between these modalities, while simultaneously learning the low-dimensional embeddings that reflect this translation. Due to the modular structure of the Flexynesis, we can attach an MLP for one or more target variables as supervisors for regression, classification, or survival tasks.

In order to demonstrate this feature, we designed an experiment using the genome-wide gene essentiality scores measured for more than 1000 cell lines as part of the DepMap project [25]. The DepMap database contains measurements of cellular proliferation after perturbation of all protein coding genes. It has been previously shown that for a given cell line, the gene expression profiles of the cell lines can be used to predict the gene essentiality scores [26,27]. Here, we carried out a similar approach, where gene expression profiles of genes across cell lines were used as input with a goal to reconstruct the cancer cell line dependency scores of the same genes. We expanded this approach to a multi-modal setup, where we used two additional data modalities besides the gene expression: 1) we used pre-trained large language models to generate protein sequence embeddings for the same genes using Prot-Trans [28] and obtained sequence embedding vectors for each gene (using the canonical protein sequences) 2) we used the structural and functional features of proteins (such as disorder profiles, evolutionary sequence conservation, secondary structures, post-translational modification sites) computed in the DescribePROT database [29]. Thus, each gene was represented by three data modalities: gene expression profiles across cell lines, protein sequence embeddings, and describeProt features. We used these modalities to reconstruct the gene-essentiality scores for each of the cell lines in the DepMap database. In addition, we attached a supervisor MLP to guide the network to predict the hubness-score of each gene in the genetic interaction networks obtained from the STRING database [30] assuming that the centrality of a gene in biological interaction networks could be a contributing factor in its essentiality for cell survival. Thus, the model is trained concurrently to predict both the gene essentiality score in a particular cell line (as a matrix), along with the gene hubness (as a vector). We trained the model on 70% of the genes and evaluated the model on the remaining 30% of the genes, by computing the average correlation of each cell line’s predicted gene dependency scores with the measured scores. Adding the protein sequence embeddings from the language models had a significant improvement on the performance of the model, while addition of DescribeProt features did not make an additional improvement over the protein language embeddings (Figure 5A), which suggests that LLM-based protein embeddings might be already capturing similar information to the features from describePROT. Using protein sequence embeddings also had a significant improvement on the prediction of the hubness scores of genes, while this time using DescribeProt features also had added benefit as well (Figure 5B).

**Figure 5:**
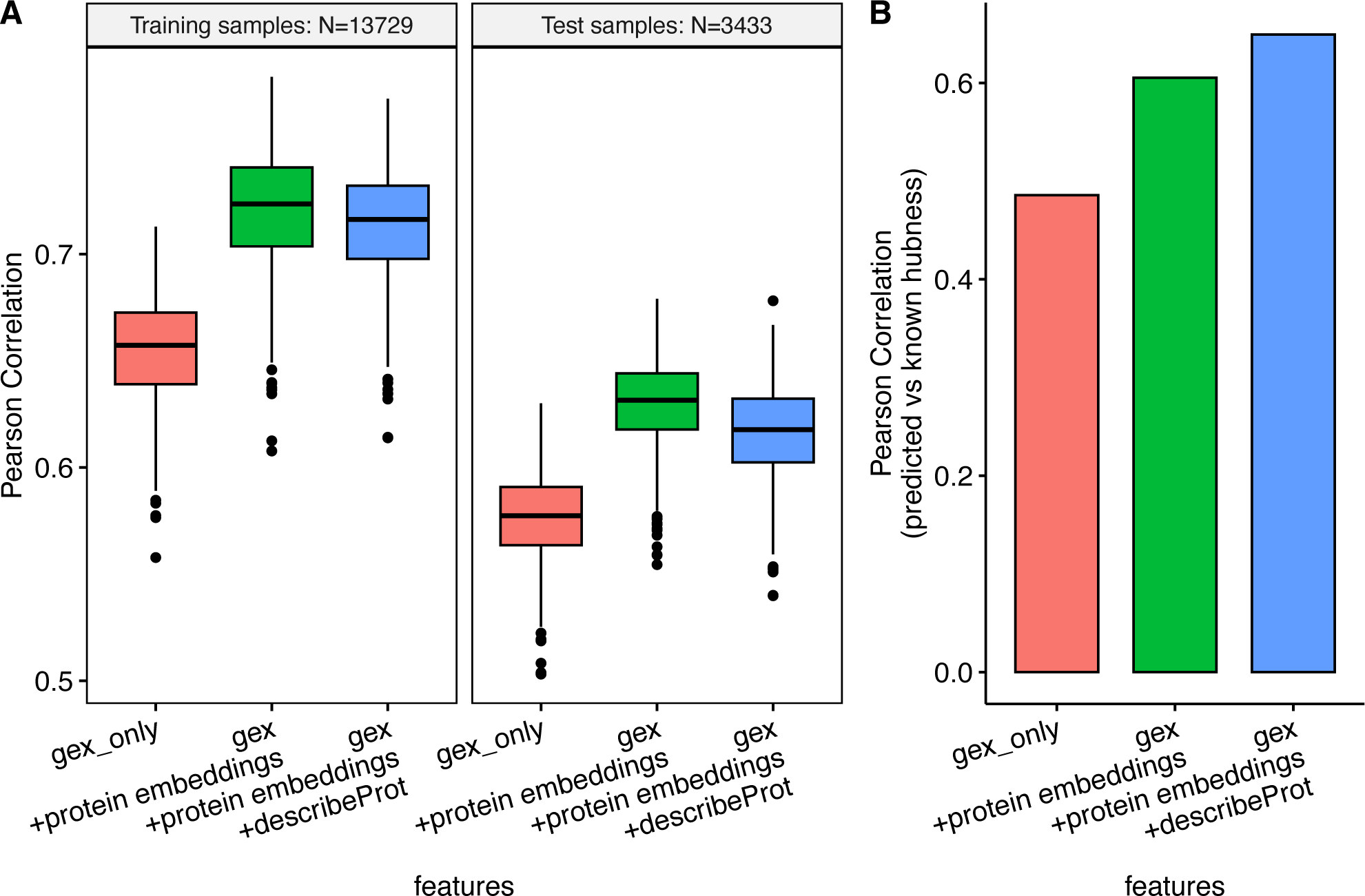
Multi-modal cross-modality prediction of gene knock-out dependency probabilities of cell lines. A cross-modal encoder-decoder model was used that takes as input a combination of input data modalities and reconstructs the CRISPR-based gene knock-out dependency probability scores of cell lines from the DepMap project. Each gene is represented with a combination of feature sets including the expression profile of the gene across cancer cell lines, Prot-Trans large language model embeddings of the gene’s canonical protein sequence, and functional/structural sequence features of the gene’s canonical protein sequence from the describePROT database. The cross-modality encoder was trained with an attached supervisor that predicts the hubness score of the gene according to its centrality in the STRING database. **Panel A)** displays the distribution of the correlation scores for each cell line’s measured gene knock-out dependency scores and the predicted scores based on different input data modality combinations: “gex_only” represents the prediction performance when using only gene expression profiles of the genes in the cell lines; “gex + protein embeddings” represents the prediction performance when using both the gene expression profiles and protein sequence embeddings from Prot-Trans; “gex + protein embeddings + describeProt” represents the prediction performance when using all three feature sets including gene expression, protein embeddings, and describeProt features. The facets in panel A represent prediction performance for the genes seen during training (N = 13729 genes) and the genes from the test set (N = 3433 genes). Panel **B)** displays the prediction performance of the hubness scores from the supervisor head that is attached to the cross-modality encoder, again using the same feature set combinations from panel A.

### Improving model performance via model fine-tuning

One of the conveniences offered by neural networks compared to classical machine learning approaches is that the neural networks trained on a source dataset can be fine-tuned on a small portion of the target dataset. This feature offers a possibility to tune the trained model on the potentially shifted distribution of the target dataset compared to the source [31]. We implemented an optional fine-tuning procedure, which uses a portion of the test dataset to modify the model parameters (following a combination of model parameter freezing strategies and different learning rates). The fine-tuned model is then evaluated on the remaining test dataset samples. In the first experiment, we trained a supervised-VAE model on seven different drug response profiles of the CCLE database and fine-tuned the trained model on different numbers of samples (100, 200, and 300 samples) from the test dataset (GDSC database). We observed that, while fine-tuning does not guarantee an improvement in model performance it can provide a boost in accuracy (Figure 6A).

**Figure 6:**
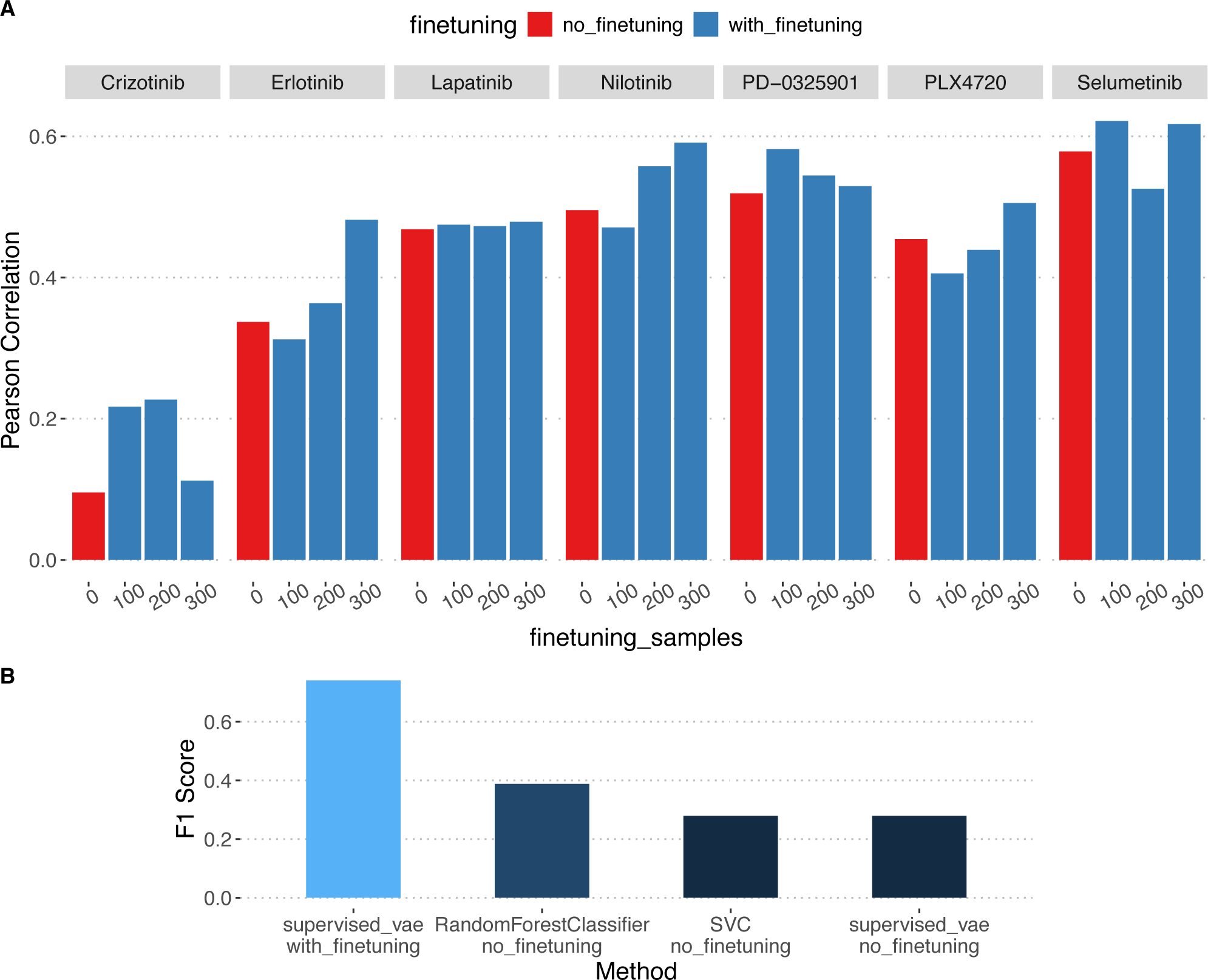
Model fine-tuning can improve prediction performance. **A)** A supervised variational auto-encoder was trained on the CCLE dataset for 7 different drug response scores and evaluated on the GDSC2 dataset. One model was trained without fine-tuning (fine-tuning = 0) and 3 different models were trained using 100, 200, and 300 samples respectively from GDSC2 for fine-tuning and the models were evaluated on the remaining unseen test samples from the GDSC2 dataset. **B)** A supervised variational autoencoder was trained on 144 Neuroblastoma patient samples from the TARGET study (using gene-expression data) to predict the MYCN gene’s amplification status. The model was evaluated on 32 neuroblastoma cell lines from the DepMap dataset for its performance in predicting the MYCN status. In the absence of fine-tuning (red bars) both the supervised variational auto-encoder and Support Vector Classifier / Random Forest Classifier predict as good as random. Using fine-tuning on 10 cell lines, the supervised variational autoencoder achieves an accuracy of 0.74 on the remaining 22 cell lines.

As the CCLE and GDSC databases have a relatively similar origin, resulting in good concordance with similar distributions, we tested fine-tuning in a separate experiment where the source (training) dataset and target (test) datasets come from completely different sources. We built models to predict the MYCN amplification status in human neuroblastoma samples from the TARGET study and use the trained model to predict the MYCN amplification status in the neuroblastoma cell lines from the DepMap database. We observed that our model performed very poorly without fine-tuning, with an F1 score of 0.3. We have verified this by evaluating the performance of a Random Forest and a Support Vector classifiers (Figure 6B). fine-tuning the model using only 10 neuroblastoma cell line samples made a significant difference in model performance, by boosting the F1 score from 0.3 to 0.75 (Figure 6B).

### Marker discovery

All model architectures, implemented in Flexynesis, are equipped with a marker discovery module based on Integrated Gradients [32,33]. In order to evaluate whether the trained Flexynesis models can capture known/expected markers, we constructed models predicting drug response, for eight drugs with known molecular targets. The models were trained on drug response data from CCLE and evaluated on the corresponding features from the GDSC dataset. We trained both a fully connected network (DirectPred) and a supervised variational auto-encoder (supervised-vae) using various data type combinations (mutations, mutations + RNA expression, and mutations + RNA expression + copy number variants). The top ten markers per drug were extracted from the best performing model among all the experiments (Figure 7A). We labeled the top markers by data type and also by the presence of the marker in civicDB [34], a database of clinically actionable genetic biomarkers of drug response. For 6 out of 8 drugs, we could find at least one known marker, present in civicDB (Figure 7B). In addition, we observe that the best performing models are never trained on single-modalities. Top markers for each of the drugs are dominated by single nucleotide variants, however, we also observe that the best performing models (Figure 7A) are the ones where the mutation data is complemented with at least the “RNA” layer, which is in line with previous findings, we and others have demonstrated before, that using the gene expression data on top of the mutation features significantly improves drug response prediction performance [35,36].

**Figure 7:**
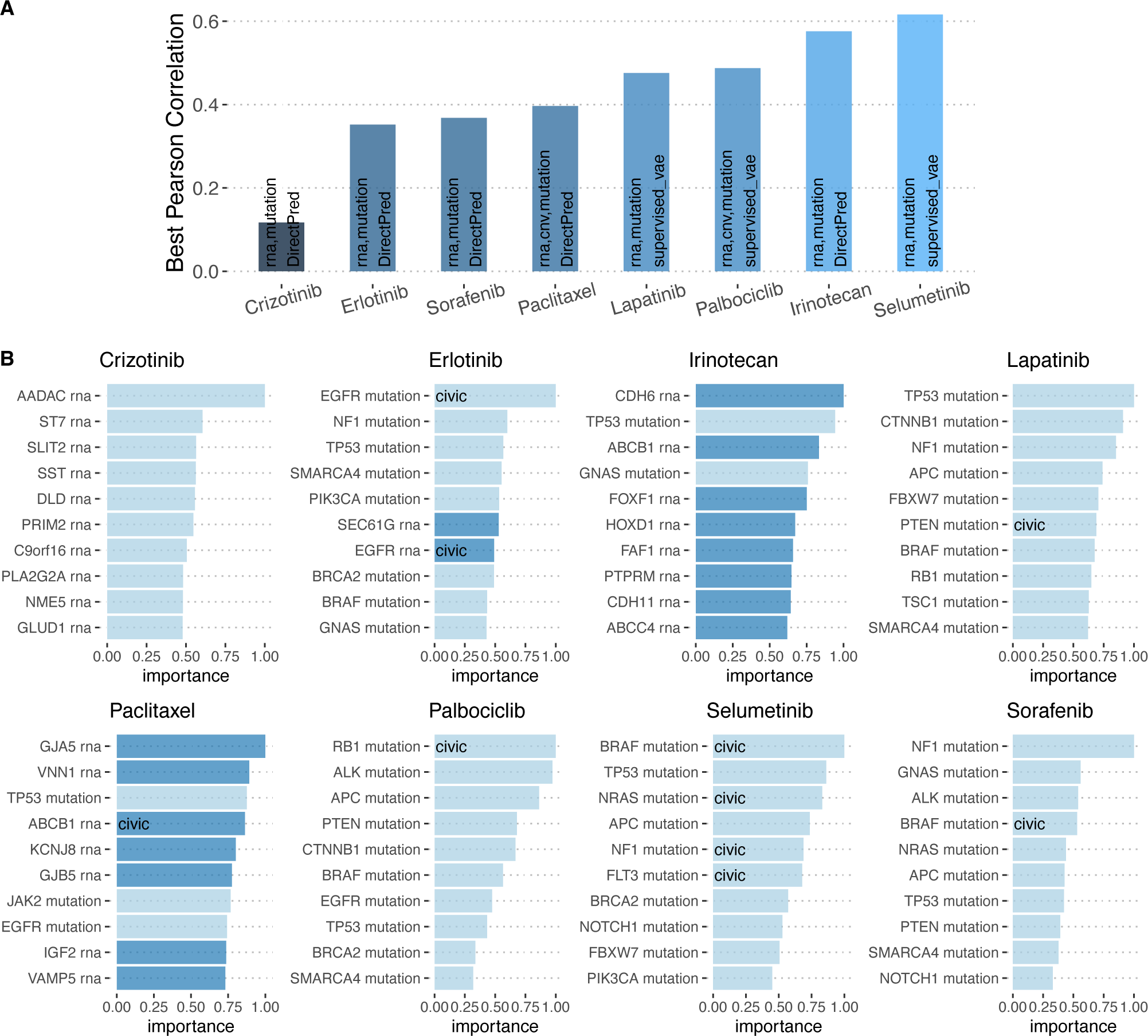
Top markers discovered using the feature importance modules of Flexynesis in predicting the drug response values (trained on the CCLE dataset and evaluated on the GDSC2 dataset). Both a fully connected network (DirectPred) and a supervised variational autoencoder (supervised vae) was trained on three combinations of data modalities: using only mutations; using mutations and RNA expression; using mutations, RNA expression, and copy number alterations. **A)** Best performing model+data type combination for each drug is displayed. **B)** The top 10 markers (in the y-axis) discovered for each drug (based on the best performing model + data type combination depicted in panel A). The markers are both labeled and colored by the corresponding data modality (dark blue: RNA expression, light blue: Mutation). The markers that are already known to be indicator markers for the corresponding drug according to the CIViC (Clinical Interpretation of Variants in Cancer) database are labeled as “civic”. The x-axis displays the relative importance of the top markers, where the best marker has a value of 1. While most drugs have dominantly mutation markers in the top 10, the best performing models always have RNA expression as an additional data modality.

### Benchmarking Pipeline

Previous benchmarking of different neural network architectures [13,14,37], showed that none of the methods outperform in all tested scenarios. It is challenging to choose the best performing neural network architecture along with the type of multi-omic modalities best suited for a given task ahead of time. Additionally, it is perfectly possible that the accuracy of the classical machine learning methods, like a random forest classifier, is sufficient for a given prediction task. Therefore, to attain the best performing model, we have to execute multiple experiments with different data type combinations, different fusion approaches, and different neural network architectures. Moreover, some tasks might benefit from building multi-task training, while others might perform better for the target variable of interest in a single-task setting.

To accommodate such combinatorial experimentations, we setup a benchmarking pipeline which can be configured to run different flavors of Flexynesis on different combinations of data modalities, different fusion options, fine-tuning options, along with a baseline performance evaluation random forests, support vector machines and random survival forests. The pipeline then builds a dashboard with rankings of different experiments in terms of prediction performances for different tasks.

We ran the benchmarking pipeline on datasets with clinically relevant outcome variables and built a dashboard of rankings of the different experiments (See <u>dashboard</u> and Supplementary Table 2). We designed 13 different tasks across 4 different datasets in a total of 480 different experiments, where we tested different tools, tool flavors, omics data type combinations, data fusion and fine-tuning options. Immediate observation confirms previous findings that no single neural network model outperforms others in all tasks. Of the 13 tasks, the top ranking method was usually a deep learning model (Figure 8A-B) however classical machine learning models SVMs and Random Forests also often perform comparably well (Supplementary Table 2). Furthermore, we compared the deep learning models in terms of omics data modality fusion options (see Methods: Data modality fusion options). Among the best performing models, the intermediate fusion shows a slightly better result than the early fusion approach, however the difference doesn’t seem significant (Figure 8C). Similarly, fine tuned deep learning models show a slight but insignificant improvement over the counterparts with no fine-tuning (Figure 8D). Finally, among the GNN models, the choice of SAGE convolution method yields slightly better results in our experiments (Figure 8E). These experiments suggest that the intermediate fusion option with fine-tuning is likely to yield better results, however, it is probably not possible to generalize to all possible situations. The value of different approaches is task specific, therefore we advise running multiple experiments to obtain the best model for the dataset at hand. This accessory pipeline ameliorates the execution of such experiments. The pipeline is available at https://github.com/BIMSBbioinfo/flexynesis-benchmarks.

**Figure 8:**
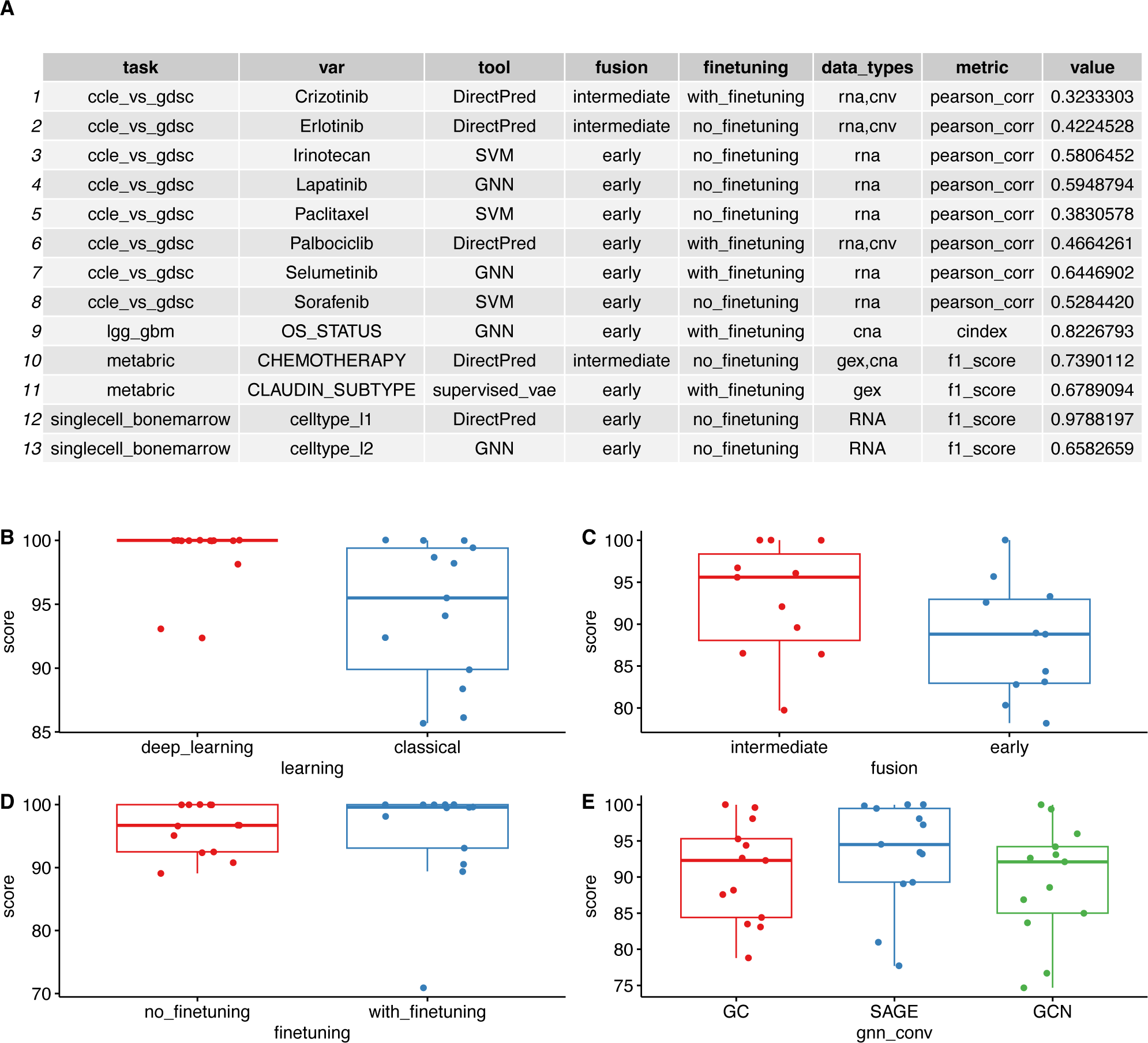
Summary of benchmarking results using different tools and data integration options with the Flexynesis benchmarking pipeline (See Supplementary Table 2 for full table of results). The “score” in the y-axis for panels B-E are scaled to 100 (where the best model gets a score of 100) in order to enable comparison of different scoring metrics. **A)** Best performing model setup for each of 13 different tasks. **B)** Comparison of the best result of among any deep learning method with the best result of the classical machine learning approaches. **C)** Comparison of the best performing data fusion strategies for the deep learning models in experiments where there were at least 2 data modalities. **D)** Comparison of the best performing model with/without the fine-tuning option. **E)** Comparison of the best performing GNN models in terms of different convolution options.

## Discussion

In this paper, we presented Flexynesis, a deep learning based bulk multi-omics integration suite with a focus on (pre-)clinical variable prediction. Despite the availability of many published deep learning-based methods, the main reason for developing this package was to provide an improved user experience when adapting deep learning for multi-omic data analysis. Existing methods lack one or more of the important components, where the absence of any of these components creates significant overhead for the users when adapting deep learning applications in their experiments. We provide a package that is easily installable, supported with good documentation, real-life benchmarking datasets with example applications, and automates data cleanup and harmonization, feature selection, hyperparameter optimisation, model evaluation, and feature importance ranking. The package is designed in a way that the user can easily switch different kinds of model architectures, can easily decide which data types to use in modeling by simply providing a list of files, experiment with modeling different kinds of clinical variables in single-task or multi-task settings and build models for supervised (regression, classification, survival), unsupervised, or cross-modality tasks without having any in-depth experience in building deep learning architectures. Thus, the user can focus on the biological context of the study and come up with interesting questions to solve with this diverse toolkit.

It has been previously shown that deep learning may struggle to outperform classical machine learning methods [15–17], which we have also observed in a subset of our benchmarking experiments. Even though classical machine learning methods might perform as good as a neural network in certain situations, the decision to use deep learning is not guided solely by the prediction performance for a given task. Deep learning offers a broader level of flexibility such as tolerance for missing labels, support for multi-task modeling, enables both supervised/unsupervised/and matrix-to-matrix predictions while simultaneously allowing dimension reduction. Furthermore, pre-trained deep learning models can be fine-tuned on a separate dataset enabling transfer learning. Finally, deep learning gains a competitive advantage with increasing amounts of data [37,38], which should be more commonplace in clinical research as multi-omic profiling becomes easier and cheaper over time.

By easily adapting Flexynesis into a bioinformatics pipeline, we have assessed both the relative performance of different flavors of deep learning architectures, along with other parametric choices one can make in multi-omics integration such as the combination of data modalities, different fusion options, and fine-tuning options. The benchmarking pipeline we built with various real-life datasets should allow both the developers in assessing the strengths and weaknesses of the novel features contributed to the package, but also guide the users to make choices based on the nature of the modeling task.

For future development of the toolkit, each of the multiple components will be easily expanded by implementing alternative methods. Currently we offer multiple alternative models, but not so many alternative algorithms for feature selection, hyperparameter optimisation, or marker discovery. We plan to implement alternative hyperparameter optimization algorithms provided by libraries such as Ray Tune [39] or Optuna [40], and expand the marker discovery, using various ranking algorithms available within the Captum library. Feature selection can be extended using unsupervised feature selection methods such as Fractal Autoencoders [41].

As a final remark, it is important to note that what we developed here is not a set of novel deep learning algorithms. None of the components we built are novel, however the innovation comes from how these components are brought together into a usable package. Flexynesis improves user experience and makes multi-omic deep learning accessible to a broader audience.

## Methods

Flexynesis is a pytorch-lightning based deep learning framework designed for bulk multi-omics data integration with a focus on precision oncology applications, however it is possible to use it for any tabular multi-modal datasets. Flexynesis workflow consists of the following main steps: importing the multi-omics data and metadata for training and testing samples, running a bayesian sequential hyperparameter optimisation routine using scikit-optimize package [42] on the training dataset and choosing the best model parameters in terms of validation metrics, evaluating the best performing model on the test (holdout) dataset, and computing the ranking of the input features in terms of importance using the Captum package [33]. If the user opts for a fine-tuning procedure, the trained model is fine-tuned on a subsample of the testing set and the fine-tuned model is evaluated on the remaining test samples.

### Importing the training and test datasets

Flexynesis expects a path to a data folder which contains training and testing data. Both training and testing data folders should contain at least one matching data modality (e.g. omics1.csv, omics2.csv … ) as a data matrix and a meta-data file that contains sample labels for each sample (clin.csv). The omics data files contain omic profiles of samples where the column names represent unique sample/patient ids and row names represent the profiled omic features. The sample metadata file (clin.csv) contains the unique sample names in rows and clinical features (outcome variables) as column names.

During the data import, Flexynesis checks for common file format errors and cross-checks information available in omics data files and metadata files to make sure that both training and testing datasets are ready for downstream analysis. After the sanity checks, the training data is further processed. Common issues with tabular data such as missing values are imputed, features with low variance are removed, samples with no available features are dropped. After the data cleanup, depending on the user’s requirements, a feature selection is implemented to keep the top most informative features based on the Laplacian Scoring method [43]. Among the top most informative features, highly redundant features are also dropped to keep unique and informative features. The feature selection is done for each data modality separately. The user can choose to keep a minimum number of features per data modality.

In case the user opted to use a graph convolutional network, the genetic interaction networks are downloaded from the STRING database (according to the requested organism id) and the training data modalities are filtered to keep only the features that are found in the interaction networks.

After feature selection, the training data is scaled and centered and optionally log-transformed. Once the modifications to the training data are finished, testing data is harmonized with the training data to make it compatible with the final model. To avoid data leakage, testing data is only scaled/centered using the scaling factors learned from the training data and the features selected for training data are kept in the testing data.

Thus the testing data does not influence feature selection or data normalization. All omic data and sample labels are finally converted into pytorch tensors.

### Hyperparameter optimisation

In the current implementation of the Flexynesis package, a Bayesian sequential hyperparameter optimization procedure is followed. Initially a random set of model specific hyperparameters are assigned and scikit-optimize package [42] is used to suggest different parameters after each hyperparameter optimization iteration. The user decides on how many iterations to carry out. The commonly optimized hyperparameters are “latent_dim”: the number of units to use for the encoding (the number of dimensions to aim for the sample embeddings) per data modality, “hidden_dim_factor”: the size of the hidden layer units in relation to the size of the previous network layer. Instead of setting this to absolute value terms, we decided to make it into relative values so that the parameter search behaves similarly depending on different input sizes, as different data modalities may have different number of features, thus having different input layer sizes in the network. “Supervisor_hidden_dim”: represents the number of units to use in the hidden layer of the MLP heads. The input layer size of this MLP is the total size of the latent factors (number of modalities x latent dim parameter). “lr”: the learning rate for the ADAM optimiser [44], “epochs”: the max number of epochs to continue the training. We use the ‘early_stop_patience’ callback so that the training is stopped if the validation loss values aren’t improving after a set number of epochs. Using the early stop patience significantly improves training speed and also avoids overfitting on the training data, thus improving model generalization on test data.

### Model/network/encoding options

Flexynesis currently contains a selection of architectures which can be used to train the models.

● **DirectPred**: A multi-task fully connected neural network for direct prediction of one or more target variables
● **GNN**: A graph neural network that by default uses the STRING database as interaction networks. Different graph convolution options are available: GraphConv [45], GCNConv [46], and SAGEConv [47]. Currently supports only early-fusion of data modalities.
● **supervised_vae**: A variational autoencoder model architecture with MMD loss.
● **MultiTripletNetwork**: A fully connected neural network implemented with a triplet loss-based contrastive learning
● **CrossModalPred**: A cross-modality encoder/predictor, which is a special implementation of the variational auto-encoders, in which the input data modalities and output data modalities can be set to different subsets of the available data modalities.

All of these networks can be augmented with one or more Multi-Layered-Perceptrons (MLPs) depending on the number of target variables the user wants to build a prediction model for. The user can select one or more target variables for regression/classification tasks. On top of these regression/classification heads, a survival MLP can be added for which the user needs to provide two variables, where time represents the time since last followup, and event a binary value (0 or 1) which represents whether an event has occurred since the last follow up. An event can be any clinically relevant event such as disease progression or a death event.

### Data modality fusion options

Flexynesis supports two kinds of data modality fusion options for fusing the omics layers. With “early fusion”, all input omic matrices are concatenated prior to training. With intermediate fusion, the input omic matrices are individually propagated through dedicated encoding networks. The output layer of the encoding networks (or the latent layer in the auto-encoder architectures) are concatenated and used as input to the MLP heads for each target/survival variable.

### Model training and loss functions

During model training, the training data is split by default into 80/20 portions for training and validation. The user can also select to do a k-fold cross-validation, in which the training data will be split into k-folds. For each MLP head dedicated to the corresponding outcome variable, a loss function is computed according to the variable types. If the variable is a continuous/numeric variable, a mean-squared-error loss is computed. If the variable is a categorical variable, a cross-entropy loss value is computed. If the variable is a survival variable, the cox-proportional hazards loss function is computed. The VAE models have an additional loss value: Maximum Mean Discrepancy (MMD) Loss [24]. The MultiTripletNetwork models use a triplet loss for contrastive learning, where the similarity between the anchor sample and positive examples are maximized, while the similarity between the anchor sample and the negative examples are minimized [48].

Depending on the model architecture and the number of MLP heads, there may be multiple loss values computed for a training task. The total loss is computed by summing up the individual loss values. However, as different loss functions can have different scales, it may be beneficial or even necessary to have a weighting schema to avoid one of the loss values to dominate the training. For this, we implemented the uncertainty weighting method [49], which can be disabled.

The total validation loss guides the training process. The final validation loss obtained from the training run is used to inform the hyperparameter optimiser to set the next set of hyperparameters for the next run.

### Model fine-tuning

When the user opts for a fine-tuning procedure, a portion of the test samples (user defined) are used to fine-tune the trained model parameters. The fine-tuning can be beneficial in cases of dramatic shifts in dataset distributions between training and test datasets. The fine-tuning procedure consists of a five-fold cross-validation scheme on a grid searching a combination of different learning rates and different model parameter freezing strategies (freeze the encoders, freeze the MLP heads or freeze none). Again an early stop callback is used with a low patience (3 epochs) to avoid overfitting to the testing dataset. The best model from this cross-validation scheme is chosen as the fine tuned model to be evaluated on the remaining test samples.

### Model performance evaluation metrics

Once a model is optimized on the training/validation sets, the model is evaluated on the testing dataset. For regression tasks, we compute the mean squared error, R-squared, and the Pearson correlation coefficients to evaluate the performance of a model. For classification tasks, we compute the balanced accuracy, F1 score, and kappa statistic. For survival tasks, we compute the Harrel’s C-index as the model evolution metric.

### Feature importance calculation for marker detection

After model training, most important features for each target variable and for each factor within the target variable are calculated using the Integrated Gradients method [32].

### Assessment of baseline performance

Flexynesis also contains functions to evaluate the prediction performance of classical machine learning algorithms on the same task. For regression and classification tasks, random forests [50] and support vector machines [51], and for survival tasks, random survival forests [52] are trained using a 5-fold cross-validation scheme where hyperparameter optimisation is carried out on the training data and best performing model is evaluated on the test dataset with the same metrics as we use for the neural network models. Scikit-learn library was extensively utilized for these methods and computing the evaluation metrics [53].

### Network Analysis

Human genetic interaction networks were downloaded from the STRING database [30] and network centrality measure (hubness score) was calculated using the igraph R package [54].

### Datasets

#### CCLE

Multi-omic and drug response data for the cell lines from the CCLE [10] was downloaded from https://zenodo.org/records/3905462 and processed using the PharmacoGx R package [55].

#### GDSC2

Multi-omic and drug response data for the cell lines from the GDSC was downloaded from https://zenodo.org/record/3905481 and processed using the PharmacoGx R package [55].

### Lower Grade Glioma (LGG) and Glioblastoma Multiforme (GBM) Merged Cohorts

The merged cohorts for LGG and GBM dataset [20] were downloaded from Cbioportal: https://www.cbioportal.org/study/summary?id=lgggbm_tcga_pub.

### METABRIC

Multi-omic data for the metastatic breast cancer cohort from the METABRIC study [22] was downloaded from Cbioportal: https://www.cbioportal.org/study/summary?id=brca_metabric

### Single-cell CITE-Seq of Bone Marrow

Single-cell CITE-Seq dataset [19] was downloaded and processed using Seurat (v5.1.0) [56]. 5000 cells were randomly sampled for training and 5000 cells were sampled for testing.

### DepMap

The omics data, CRISPR screens and PRISM drug screening data for cell lines from the DepMap project [25] was downloaded from the DepMap Portal (https://DepMap.org/portal).

### TARGET Neuroblastoma

Neuroblastoma patient cohort from the TARGET study was downloaded from the Cbioportal (https://www.cbioportal.org/study/summary?id=nbl_target_2018_pub).

### TCGA Data

The TCGA datasets were downloaded using the TCGABiolinks package [57].

### Prot-Trans Sequence Embeddings

Protein sequence embeddings for each gene was obtained using the prot_t5_xl_uniref50 transformer model (available at https://huggingface.co/Rostlab/prot_t5_xl_uniref50) [28]. The canonical protein sequences of the human proteome were downloaded from the UniProt database [58]. For each protein sequence, the transformer model outputs a numeric matrix of the dimensions 1024 x N where N is the number of amino-acids in the protein sequence. For each protein sequence, the row-wise averages were calculated to obtain protein-level 1024 dimensional vector embeddings.

### describePROT

Structural/functional features of human protein sequences were downloaded from the describePROT database [29]:

http://biomine.cs.vcu.edu/servers/DESCRIBEPROT/download_database_value/9606_v alue.csv.

### Data visualization

Data visualization methods implemented in the Flexynesis package uses Matplotlib [59] and Seaborn [60] and lifelines [61] python libraries.

We used icons from www.flaticon.com in the graphical abstract of this manuscript.

### Clustering

Flexynesis is equipped with utility functions to cluster a given matrix and choose optimal clusters. Currently two clustering methods are supported: Louvain clustering from the community package [62] and k-means algorithm from the scikit-learn package [53], where the clustering can be done for different values of k and the optimal clustering result can be selected by Silhouette score rankings.

### Data and code availability

Flexynesis software package is available at: https://github.com/BIMSBbioinfo/flexynesis and https://pypi.org/project/flexynesis/. The accessory benchmarking pipeline utilizing Flexynesis is available at: https://github.com/BIMSBbioinfo/flexynesis-benchmarks.

## Author contributions

B.U. conceived the project, implemented the Flexynesis software and the benchmarking pipeline, designed the experiments, carried out data analyses, wrote the paper, and supervised the software contributors. T.S. contributed to the GNN model architecture. A.S. helped with adding different graph convolution options. R.W. helped package the software for pypi and Guix. M.M.S. designed the dashboard of the benchmarking pipeline results. V.F. contributed to the conception of the project, helped with interpretation of results, and edited the manuscript. A.A. contributed to the conception of the project, helped with data interpretation and designing experiments, edited the manuscript and provided funding and resources.

## Acknowledgements

We are thankful to Aaron Kollotzek, Alexander Blume, Artur Manukyan, Fabian Janosch Krueger, Jacqueline Jansen, and Jonas Freimuth for joining the user experience hackathon and testing the software. We also thank Dan Munteanu and Martin Siegert for their help with the compute servers.

The results published here are in whole or part based upon data generated by the TCGA Research Network: https://www.cancer.gov/tcga.

The results published here are in whole or part based upon data generated by the Therapeutically Applicable Research to Generate Effective Treatments (https://www.cancer.gov/ccg/research/genome-sequencing/target) initiative, phs000218. The data used for this analysis are available at the Genomic Data Commons (https://portal.gdc.cancer.gov).

## References

1. Sung H, Ferlay J, Siegel RL, Laversanne M, Soerjomataram I, Jemal A, et al. Global Cancer Statistics 2020: GLOBOCAN Estimates of Incidence and Mortality Worldwide for 36 Cancers in 185 Countries. CA Cancer J Clin. 2021;71. doi:10.3322/caac.21660

2. Hanahan D, Weinberg RA. Hallmarks of cancer: the next generation. Cell. 2011;144. doi:10.1016/j.cell.2011.02.013

3. Hasin Y, Seldin M, Lusis A. Multi-omics approaches to disease. Genome Biol. 2017;18. doi:10.1186/s13059-017-1215-1

4. Worthey EA, Mayer AN, Syverson GD, Helbling D, Bonacci BB, Decker B, et al. Making a definitive diagnosis: successful clinical application of whole exome sequencing in a child with intractable inflammatory bowel disease. Genet Med. 2011;13. doi:10.1097/GIM.0b013e3182088158

5. Joshi A, Rienks M, Theofilatos K, Mayr M. Systems biology in cardiovascular disease: a multiomics approach. Nat Rev Cardiol. 2020;18: 313–330.

6. Karczewski KJ, Snyder MP. Integrative omics for health and disease. Nat Rev Genet. 2018;19: 299–310.

7. Guan F, Ni T, Zhu W, Williams LK, Cui L-B, Li M, et al. Integrative omics of schizophrenia: from genetic determinants to clinical classification and risk prediction. Mol Psychiatry. 2021;27: 113–126.

8. Chen R, Mias GI, Li-Pook-Than J, Jiang L, Lam HYK, Chen R, et al. Personal Omics Profiling Reveals Dynamic Molecular and Medical Phenotypes. Cell. 2012;148: 1293–1307.

9. Park YH, Im SA, Park K, Wen J, Lee KH, Choi YL, et al. Longitudinal multi-omics study of palbociclib resistance in HR-positive/HER2-negative metastatic breast cancer. Genome Med. 2023;15. doi:10.1186/s13073-023-01201-7

10. Barretina J, Caponigro G, Stransky N, Venkatesan K, Margolin AA, Kim S, et al. The Cancer Cell Line Encyclopedia enables predictive modelling of anticancer drug sensitivity. Nature. 2012;483: 603–607.

11. Steyaert S, Pizurica M, Nagaraj D, Khandelwal P, Hernandez-Boussard T, Gentles AJ, et al. Multimodal data fusion for cancer biomarker discovery with deep learning. Nature Machine Intelligence. 2023;5: 351–362.

12. Bersanelli M, Mosca E, Remondini D, Giampieri E, Sala C, Castellani G, et al. Methods for the integration of multi-omics data: mathematical aspects. BMC Bioinformatics. 2016;17: 167–177.

13. Hauptmann T, Kramer S. A fair experimental comparison of neural network architectures for latent representations of multi-omics for drug response prediction. BMC Bioinformatics. 2023;24: 1–25.

14. Leng D, Zheng L, Wen Y, Zhang Y, Wu L, Wang J, et al. A benchmark study of deep learning-based multi-omics data fusion methods for cancer. Genome Biol. 2022;23: 1–32.

15. Grinsztajn L, Oyallon E, Varoquaux G. Why do tree-based models still outperform deep learning on tabular data? 2022. Available: http://arxiv.org/abs/2207.08815

16. Dong Y, Zhou S, Xing L, Chen Y, Ren Z, Dong Y, et al. Deep learning methods may not outperform other machine learning methods on analyzing genomic studies. Front Genet. 2022;13. doi:10.3389/fgene.2022.992070

17. Shwartz-Ziv R, Armon A. Tabular Data: Deep Learning is Not All You Need. 2021. Available: http://arxiv.org/abs/2106.03253

18. Yang W, Soares J, Greninger P, Edelman EJ, Lightfoot H, Forbes S, et al. Genomics of Drug Sensitivity in Cancer (GDSC): a resource for therapeutic biomarker discovery in cancer cells. Nucleic Acids Res. 2013;41. doi:10.1093/nar/gks1111

19. Stuart T, Butler A, Hoffman P, Hafemeister C, Papalexi E, Mauck WM, et al. Comprehensive Integration of Single-Cell Data. Cell. 2019;177: 1888–1902.e21.

20. Ceccarelli M, Barthel FP, Malta TM, Sabedot TS, Salama SR, Murray BA, et al. Molecular Profiling Reveals Biologically Discrete Subsets and Pathways of Progression in Diffuse Glioma. Cell. 2016;164: 550–563.

21. Katzman JL, Shaham U, Cloninger A, Bates J, Jiang T, Kluger Y. DeepSurv: personalized treatment recommender system using a Cox proportional hazards deep neural network. BMC Med Res Methodol. 2018;18: 1–12.

22. Curtis C, Shah SP, Chin S-F, Turashvili G, Rueda OM, Dunning MJ, et al. The genomic and transcriptomic architecture of 2,000 breast tumours reveals novel subgroups. Nature. 2012;486: 346–352.

23. Koschmann C, Calinescu A-A, Nunez FJ, Mackay A, Fazal-Salom J, Thomas D, et al. ATRX loss promotes tumor growth and impairs nonhomologous end joining DNA repair in glioma. Sci Transl Med. 2016;8: 328ra28.

24. Zhao S, Song J, Ermon S. InfoVAE: Information Maximizing Variational Autoencoders. 2017. Available: http://arxiv.org/abs/1706.02262

25. DepMap B. DepMap 24Q2 Public. Figshare+; 2024. doi:10.25452/figshare.plus.25880521.v1

26. Itzhacky N, Sharan R. Prediction of cancer dependencies from expression data using deep learning. Mol Omics. 2021;17: 66–71.

27. Rosenski J, Shifman S, Kaplan T. Predicting gene knockout effects from expression data. BMC Med Genomics. 2023;16: 1–13.

28. Elnaggar A, Heinzinger M, Dallago C, Rehawi G, Wang Y, Jones L, et al. ProtTrans: Toward Understanding the Language of Life Through Self-Supervised Learning. [cited 8 Jul 2024]. Available: https://ieeexplore.ieee.org/document/9477085

29. Zhao B, Katuwawala A, Oldfield CJ, Dunker AK, Faraggi E, Gsponer J, et al. DescribePROT: database of amino acid-level protein structure and function predictions. Nucleic Acids Res. 2021;49: D298–D308.

30. Szklarczyk D, Kirsch R, Koutrouli M, Nastou K, Mehryary F, Hachilif R, et al. The STRING database in 2023: protein–protein association networks and functional enrichment analyses for any sequenced genome of interest. Nucleic Acids Res. 2022;51: D638–D646.

31. Kouw WM, Loog M. An introduction to domain adaptation and transfer learning. 2018. Available: http://arxiv.org/abs/1812.11806

32. Sundararajan M, Taly A, Yan Q. Axiomatic Attribution for Deep Networks. 2017. Available: http://arxiv.org/abs/1703.01365

33. Kokhlikyan N, Miglani V, Martin M, Wang E, Alsallakh B, Reynolds J, et al. Captum: A unified and generic model interpretability library for PyTorch. 2020. Available: http://arxiv.org/abs/2009.07896

34. Griffith M, Spies NC, Krysiak K, McMichael JF, Coffman AC, Danos AM, et al. CIViC is a community knowledgebase for expert crowdsourcing the clinical interpretation of variants in cancer. Nat Genet. 2017;49: 170–174.

35. Baranovskii A, Gündüz IB, Franke V, Uyar B, Akalin A. Multi-Omics Alleviates the Limitations of Panel Sequencing for Cancer Drug Response Prediction. Cancers . 2022;14. doi:10.3390/cancers14225604

36. Malik V, Kalakoti Y, Sundar D. Deep learning assisted multi-omics integration for survival and drug-response prediction in breast cancer. BMC Genomics. 2021;22: 1–11.

37. Hanczar B, Bourgeais V, Zehraoui F. Assessment of deep learning and transfer learning for cancer prediction based on gene expression data. BMC Bioinformatics. 2022;23: 1–23.

38. Goodfellow I, Bengio Y, Courville A. Deep Learning. MIT Press; 2016.

39. Liaw R, Liang E, Nishihara R, Moritz P, Gonzalez JE, Stoica I. Tune: A Research Platform for Distributed Model Selection and Training. 2018. Available: http://arxiv.org/abs/1807.05118

40. Akiba T, Sano S, Yanase T, Ohta T, Koyama M. Optuna: A Next-generation Hyperparameter Optimization Framework. 2019. Available: http://arxiv.org/abs/1907.10902

41. Wu X, Cheng Q. Fractal Autoencoders for Feature Selection. 2020. Available: http://arxiv.org/abs/2010.09430

42. scikit-optimize/scikit-optimize: v0.5.2. [cited 8 Jul 2024]. doi:10.5281/zenodo.1207017

43. He X, Cai D, Niyogi P. Laplacian Score for Feature Selection. Adv Neural Inf Process Syst. 2005;18. Available: https://proceedings.neurips.cc/paper_files/paper/2005/file/b5b03f06271f8917685d14cea7c6c50a-Paper.pdf

44. Kingma DP, Ba J. Adam: A Method for Stochastic Optimization. 2014. Available: http://arxiv.org/abs/1412.6980

45. Morris C, Ritzert M, Fey M, Hamilton WL, Lenssen JE, Rattan G, et al. Weisfeiler and Leman Go Neural: Higher-order Graph Neural Networks. 2018. Available: http://arxiv.org/abs/1810.02244

46. Kipf TN, Welling M. Semi-Supervised Classification with Graph Convolutional Networks. 2016. Available: http://arxiv.org/abs/1609.02907

47. Hamilton WL, Ying R, Leskovec J. Inductive Representation Learning on Large Graphs. 2017. Available: http://arxiv.org/abs/1706.02216

48. Hoffer E, Ailon N. Deep metric learning using Triplet network. 2014. Available: http://arxiv.org/abs/1412.6622

49. Kendall A, Gal Y, Cipolla R. Multi-Task Learning Using Uncertainty to Weigh Losses for Scene Geometry and Semantics. 2017. Available: http://arxiv.org/abs/1705.07115

50. Breiman L. Random Forests. Mach Learn. 2001;45: 5–32.

51. Cortes C, Vapnik V. Support-vector networks. Mach Learn. 1995;20: 273–297.

52. Ishwaran H, Kogalur UB, Blackstone EH, Lauer MS. Random survival forests. 2008 [cited 9 Jul 2024]. doi:10.1214/08-AOAS169

53. Pedregosa F, Varoquaux G, Gramfort A, Michel V, Thirion B, Grisel O, et al. Scikit-learn: Machine Learning in Python. J Mach Learn Res. 2011;12: 2825–2830.

54. Antonov M, Csárdi G, Horvát S, Müller K, Nepusz T, Noom D, et al. igraph enables fast and robust network analysis across programming languages. 2023. Available: http://arxiv.org/abs/2311.10260

55. Smirnov P, Safikhani Z, El-Hachem N, Wang D, She A, Olsen C, et al. PharmacoGx: an R package for analysis of large pharmacogenomic datasets. Bioinformatics. 2015;32: 1244–1246.

56. Hao Y, Hao S, Andersen-Nissen E, Mauck WM, Zheng S, Butler A, et al. Integrated analysis of multimodal single-cell data. Cell. 2021;184. doi:10.1016/j.cell.2021.04.048

57. Colaprico A, Silva TC, Olsen C, Garofano L, Cava C, Garolini D, et al. TCGAbiolinks: an R/Bioconductor package for integrative analysis of TCGA data. Nucleic Acids Res. 2015;44: e71–e71.

58. The UniProt Consortium, Bateman A, Martin M-J, Orchard S, Magrane M, Ahmad S, et al. UniProt: the Universal Protein Knowledgebase in 2023. Nucleic Acids Res. 2022;51: D523–D531.

59. Hunter JD. Matplotlib: A 2D Graphics Environment. Comput Sci Eng. 2007;9: 90–95.

60. Waskom ML. seaborn: statistical data visualization. Journal of Open Source Software. 2021;6: 3021.

61. Davidson-Pilon C. lifelines: survival analysis in Python. J Open Source Softw. 2019;4: 1317.

62. Blondel VD, Guillaume J-L, Lambiotte R, Lefebvre E. Fast unfolding of communities in large networks. J Stat Mech. 2008;2008: P10008.

